# Identification of Interleukin1β as an Amplifier of Interferon alpha-induced Antiviral Responses

**DOI:** 10.1101/2020.03.10.985390

**Authors:** Katharina Robichon, Tim Maiwald, Marcel Schilling, Annette Schneider, Joschka Willemsen, Florian Salopiata, Melissa Teusel, Salopiata, Clemens Kreutz, Christian Ehlting, Jun Huang, Sajib Chakraborty, Xiaoyun Huang, Georg Damm, Daniel Seehofer, Philipp A. Lang, Johannes G. Bode, Marco Binder, Ralf Bartenschlager, Jens Timmer, Ursula Klingmüller

## Abstract

The induction of an interferon-mediated response is the first line of defense against pathogens such as viruses. Yet, the dynamics and extent of interferon alpha (IFNα)-induced antiviral genes vary remarkably and comprise three expression clusters: early, intermediate and late. By mathematical modeling based on time-resolved quantitative data, we identified mRNA stability as well as a negative regulatory loop as key mechanisms endogenously controlling the expression dynamics of IFNα-induced antiviral genes in hepatocytes. Guided by the mathematical model, we uncovered that this regulatory loop is mediated by the transcription factor IRF2 and showed that knock-down of IRF2 results in enhanced expression of early, intermediate and late IFNα-induced antiviral genes. Co-stimulation experiments with different pro-inflammatory cytokines revealed that this amplified expression dynamics of the early, intermediate and late IFNα-induced antiviral genes can be mimicked by co-application of IFNα and interleukin1 beta (IL1β). Consistently, we found that IL1β enhances IFNα-mediated repression of viral replication. Conversely, we observed that in IL1β receptor knock-out mice replication of viruses sensitive to IFN is increased. Thus, IL1β is capable to potentiate IFNα-induced antiviral responses and could be exploited to improve antiviral therapies.

**Author Summary:** Innate immune responses contribute to the control of viral infections and the induction of interferon alpha (IFNα)-mediated antiviral responses is an important component. However, IFNα induces a multitude of antiviral response genes and the expression dynamics of these genes can be classified as early, intermediate and late. Here we show, based on a mathematical modeling approach, that mRNA stability as well as the negative regulator IRF2 control the expression dynamics of IFNα-induced antiviral genes. Knock-down of IRF2 resulted in the amplified IFNα-mediated induction of the antiviral genes and this amplified expression of antiviral genes could be mimicked by co-stimulation with IFNα and IL1β. We observed that co-stimulation with IFNα and IL1β enhanced the repression of virus replication and that knock-out of the IL1 receptor in mice resulted in increased replication of a virus sensitive to IFNα. In sum, our studies identified IL1β as an important amplifier of IFNα-induced antiviral responses.

## Introduction

Cytokines such as interferons (IFNs) are important regulators of the innate immune system, the first line of defense against microbial infection. IFNs induce in a highly dynamic process the expression of several classes of IFN-stimulated genes. The encoded proteins of these genes fulfill a variety of tasks including the clearance of viruses. To ensure effectiveness of the response and to prevent damage, the process has to be tightly controlled, which is achieved through several positive and negative feedback loops [1]. Due to the non-linearity of the underlying reactions the impact of alterations on a potential outcome is difficult to predict. IFNs such as interferon alpha (IFNα) are widely applied therapeutic agents and therefore strategies to strengthen IFN-induced responses are of major interest. However, this requires a more quantitative understanding of the interrelations between the IFN signaling pathway components and the expression of IFN-stimulated genes (ISGs) as well as insights into mechanisms shaping the response to IFNs.

A well-studied IFN-induced response is the antiviral response elicited for example by major hepatotropic RNA viruses such as the human pathogen hepatitis C virus (HCV) and the murine pathogen lymphocytic choriomeningitis virus (LCMV). Upon infection, the viral RNA is sensed by specific cellular pattern recognition receptors (PRR) that trigger the expression of interferons (IFNs) and induce expression of antiviral genes as first line of defense [2]. However, viruses can evade the antiviral response by antagonizing the induction of the effector pathways of the IFN system and establish a persistent infection. Therefore, it would be highly beneficial to identify mechanisms to enhance the IFN-induced antiviral response to reduce virus spread and improve viral clearance.

The major signal transduction pathway activated in response to type I IFNs such as IFNα is the JAK/STAT pathway [3]. Regulation of the dynamics of the JAK/STAT pathway activation and the expression of IFN-stimulated genes are important to mount an effective IFN response and to maintain cellular homeostasis. The IFNα-induced signaling pathway comprises complex negative feedback loops consisting of suppressor of cytokine signaling 1 (SOCS1) and ubiquitin-specific peptidase 18 (USP18) that jointly determine signal attenuation. In contrast, interferon regulatory factor 9 (IRF9) acts as a positive regulator of IFNα signaling. By dynamic pathway modeling it was shown that an upregulation of IRF9 can enhance the expression of ISGs [4]. Further, it was shown that the extent and duration of the expression of antiviral genes positively correlates with a reduced virus load [5] and the specific expression profiles of antiviral genes appear to be critical for shifting the balance from viral persistence to viral clearance. Therefore, the modulation of feedback loops might be harnessed to increase and prolong the duration of the IFN response and thereby contribute to improved viral clearance.

IFNα was not only shown to activate the classical JAK-STAT1 pathway, but recent publications have also reported an activation of STAT3 after IFNα treatment [6]. For example, Su et al. showed a phosphorylation of STAT3 after IFNα treatment in RAMOS cells [7] and IFNα treatment led to an increase of STAT3 phosphorylation in primary healthy dendritic cells [8] as well as B cells [9]. The activation of the different STAT molecules may promote the formation of different hetero- and homodimer pairs, resulting in different expression of the ISGs.

In addition to type I IFNs, pro-inflammatory cytokines such as interleukin 6 (IL6), interleukin-1beta (IL1β) and IFN gamma (IFNγ) [10] can contribute to the activation of an anti-microbial response. Binding of IL1β to the type I IL1 receptor (IL1R1) that is expressed on different cell types including hepatocytes results in the activation of different downstream signaling pathways. While the main pathways activated by IL1β are p38 and NFκB [11], there is evidence that IL1β can also activate STAT3 [12]. IL1β was reported to induce the protein-protein interaction between STAT3 and NFκB in hepatocytes as well as DNA binding of this complex [13], which might be involved in facilitating the recently reported NFκB-assisted DNA loading of STAT3 during the acute phase response [14]. An interplay between IFNα and IL1β has been observed previously. On the one hand, in liver samples of chronic hepatitis C patients elevated levels of IFNβ and IL1β were observed [15]. On the other hand, it was reported that IFNα and IFNβ suppress IL1β maturation in bone marrow-derived macrophages [16] and that IL1β limits excessive type I IFN production through the induction of eicosanoids [17]. Co-treatment with IFNα and IL1 β resulted in higher and more sustained STAT1 phosphorylation in Huh7 cells [18]. Thus, the physiological relevance and the underlying mechanism of a potential cross-talk between type I IFN-induced signaling and IL1β remains unknown.

Here we employ a systems biology approach that combines time-resolved quantitative experimental data and mathematical modeling. We show that mRNA stability as well as IRF2 as a negative feedback loop critically shape the distinct expression dynamics of the early, intermediate and late IFNα-induced genes. Importantly, we uncover that IL1β is capable to mimic the impact of knockdown of IRF2 and boosts the IFNα-induced antiviral gene response.

## Results

### Distinct dynamics of IFNα-induced gene expression

To characterize the temporal response induced by IFNα stimulation and to classify the induced genes based on their expression dynamics, we took advantage of our previously reported microarray analysis monitoring IFNα-induced gene expression over 24 hours in the human hepatoma cell line Huh7.5 stimulated with IFNα [4]. We used Huh7.5 cells as a model system, because this cell line has been widely used to investigate the replication of hepatotropic viruses. Utilizing these data, we focused our analysis on genes that exhibited significant upregulation (p<0.05 and average fold-change>2) in response to IFNα treatment (Fig 1A). Sorting the 53 significantly upregulated genes by the time point of maximal induction revealed three expression clusters: early, intermediate and late (Fig 1B). 21 genes classified as early were rapidly induced with a peak of maximal activation (vertical red line) one to four hours after stimulation and rapidly declined thereafter. 27 genes grouped in the intermediate cluster reached their maximal expression at six to eight hours, followed by a moderate decline. Five genes were induced late and exhibited persistent upregulation with maximal expression at 12 hours or later.

**Fig 1:**
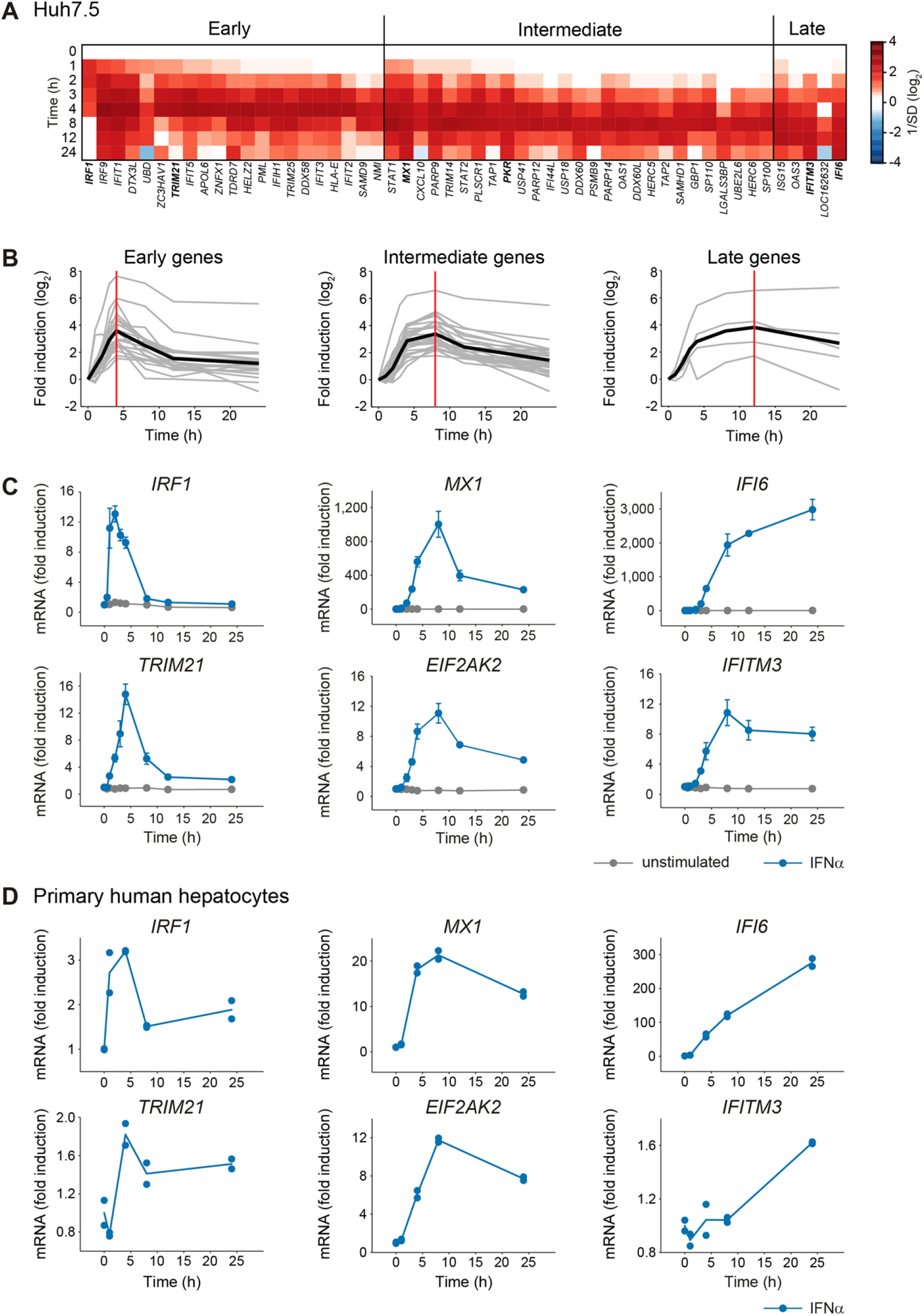
Early, intermediate and late expression profiles of IFNα-induced genes. (**A**) Microarray expression data for Huh7.5 cells stimulated with 500 U/ml IFNα. The heatmap shows the temporal expression patterns of 53 significantly upregulated genes, grouped according to their peak expression time. (**B**) The induction of the genes depicted in **A** is displayed in a time-resolved manner according to the respective groups (grey curves). The average expression of each group is indicated by a solid black line. The vertical red lines indicate the time points of maximal induction. (**C**) Huh7.5 cells were stimulated with 500 U/ml IFNα or left untreated and two representative antiviral genes per group were analyzed by qRT-PCR. The error bars represent SD of biological triplicates. (**D**) IFNα-induced mRNA expression in primary human hepatocytes. Primary human hepatocytes were growth factor depleted and stimulated with 500 U/ml IFNα. RNA was extracted at the indicated time points and analyzed using qRT-PCR. Points: experimental data; lines: average of biological duplicates.

As representatives for further analysis we selected two IFNα-induced genes with known antiviral acitivity from each group [19, 20]: IFN regulatory factor 1 (*IRF1*) and tripartite motif containing 21 (*TRIM21*) from the early group, MX dynamin-like GTPase 1 (*MX1*) and eukaryotic translation initiation factor 2-alpha kinase 2 (*EIF2AK2/PKR*) from the intermediate group, and IFNα-inducible protein 6 (*IFI6*) and IFN-induced transmembrane protein 3 (*IFITM3*) as examples from the late group. The characteristic dynamics of the IFNα-induced expression levels of each of the selected antiviral genes were verified by qRT-PCR analysis and confirmed the grouping into the early, intermediate and late cluster (Fig 1C).

To interrogate whether this dynamic behavior of IFNα-induced antiviral genes is characteristic for Huh7.5 cells and hence potentially determined by the cancer cell context or whether it is conserved in primary hepatocytes, we examined the IFNα-induced expression of the selected IFNα-induced antiviral genes in primary human hepatocytes isolated from multiple donors. Overall the observed fold change of the expression of the IFNα-induced antiviral genes was lower in primary human hepatocytes compared to Huh7.5 cells. But in line with our previous results, the anticipated dynamic behavior was observed for each of the genes tested: The early genes *IRF1* and *TRIM21* showed maximal expression between 1 and 4 hours after IFNα treatment and rapidly declined thereafter, the intermediate genes *MX1* and *EIF2AK2* showed maximal expression between six to eight hours and rather sustained expression and the late genes *IFI6* and *IFITM3* exhibited a persistent increase for the entire observation time of up to 24 hours (Fig 1D). The conserved dynamic behavior of IFNα-induced antiviral genes in Huh7.5 cells and primary human hepatocytes suggested that the expression dynamics of IFNα-induced antiviral genes is regulated by robust mechanisms maintained in hepatocytes.

### Distinct mRNA stability affects expression profiles of IFNα-induced genes

To elucidate key mechanisms that contribute to the three distinct expression profiles of the IFNα-induced antiviral genes, we first tested whether the IFNα dose-dependency differed between these groups. Comparing the half-maximal effective IFNα dose (EC_50_) of the selected IFNα-induced antiviral genes however showed that the EC_50_ of these genes ranged from 100 ± 9 to 171 ± 23 U/ml INFα and did not reveal substantial differences between the three groups (Fig 2A). Therefore, we next assessed whether the distinct expression dynamics resulted from differences in the stability of the mRNAs. To determine the mRNA half-lives of the selected IFNα-induced genes, we inhibited *de novo* transcription using actinomycin D. As shown in Fig 2B, the mRNA concentration of each of the examined antiviral genes decreased over time. To calculate the half-lives of the different mRNAs, a three-parameter exponential-decay regression was performed with the mRNA expression data. Interestingly, the mRNA expression profiles of the selected genes representing the three groups were well reflected by their mRNA half-lives (Fig 2B): mRNAs that exhibited an early-type expression profile displayed a short half-life of 30 minutes to 2 hours; intermediate-type mRNA expression showed a half-life of approximately 5 to 7 hours; and genes with sustained-type expression profiles exhibited stable mRNAs over the entire observation period.

**Fig 2:**
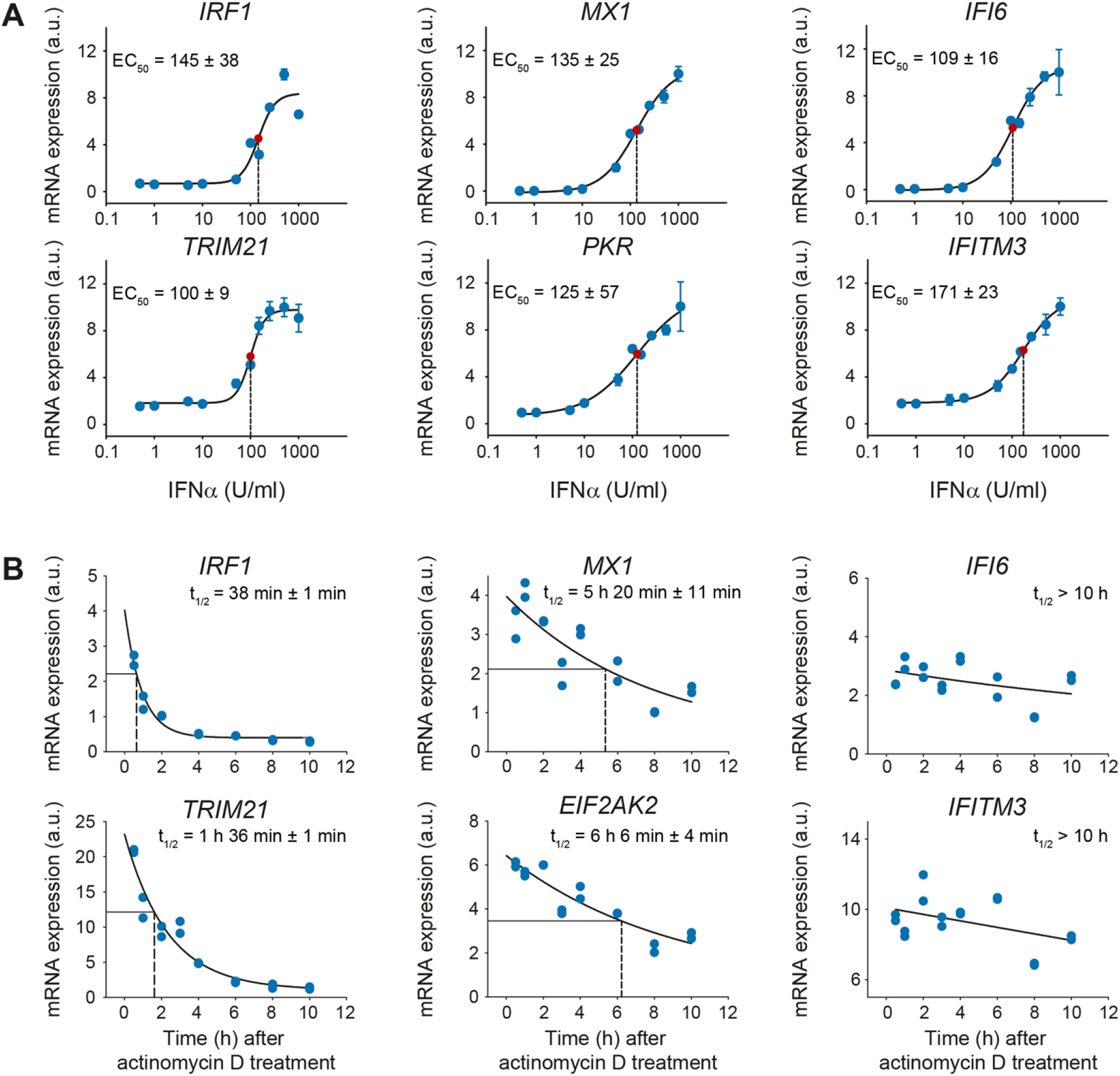
Difference in mRNA stability despite comparable dose-dependency of the expression profiles of given antiviral genes. (**A**) IFNα dose-dependent mRNA expression of antiviral genes. Huh7.5 cells were treated with increasing doses of IFNα for 4 hours. The cells were lysed, total RNA was extracted and analyzed by qRT-PCR. The error bars represent standard deviations (SD) based on biological triplicates. Regression line: sigmoidal four-parameter Hill function; red point: inflection point; dashed line: calculated EC50; a.u.: arbitrary units. (**B**) Quantification of the mRNA half-lives of the selected antiviral genes. Huh7.5 cells were stimulated with 500 U/ml IFNα for 8 hours and then treated with 5 ng/ml actinomycin D for the indicated times. Total RNA was extracted and analyzed by qRT-PCR. The data points represent biological duplicates. Regression line: three-parameter exponential decay function, dashed line: calculated RNA half-life.

Thus, the three expression groups of IFNα-induced antiviral genes did not differ in their IFNα dose dependency, but were characterized by differences in mRNA stability. However, the distinct mRNA stabilities of the three groups did not explain e.g. the observed differences in the time to maximal expression of the antiviral genes. Therefore, we concluded that additional mechanisms such as feedback loops shape the expression profiles of IFNα-induced antiviral genes.

### Analysis of the pathway structure using a dynamic model of IFNα-induced signaling

To elucidate the potential impact of feedback loops regulating the dynamic properties of the expression of IFNα-induced antiviral genes, an ordinary differential equation (ODE) model (core model) was developed (S1A Fig). The core model was based on our previously published mathematical model [4] that was expanded by introducing mRNA expression of the negative regulators SOCS1 and USP18 and the selected IFNα-induced antiviral genes. The mathematical model was calibrated based on previously published [4] and new experimental data on the activation of the JAK/STAT pathway and IFNα-induced expression of antiviral genes that were acquired for up to 24 hours post IFNα stimulation. The initial concentrations of the main pathway components were experimentally determined (S1 Table). In addition, the experimentally determined mRNA half-life values were incorporated by introducing an mRNA-specific degradation parameter for each individual mRNA.

The simulations of the core model for the IFNα-induced signaling components (exemplarily shown for phosphorylation of JAK1 and STAT1), for the induction of the positive regulator IRF9 and for the negative regulator USP18 were consistent with the experimental data (S1B Fig). However, the trajectories of the core model were not able to reproduce the induction kinetics of the early (*IRF1* and *TRIM21*, S1C Fig) and late genes (*IFI6*, S1C Fig) as well as of the negative regulatory signaling protein SOCS1 (S1B Fig). Further, the core model failed to sufficiently reproduce the downregulation of the intermediate genes *MX1* and *EIF2AK2* indicating a missing interaction (S1C Fig). Thus, we aimed to identify missing components in our mathematical model.

### IRF2 constitutes an intracellular feedback loop that negatively regulates expression of early IFNα-induced genes

To improve the capacity of the model to represent the experimental data, we incorporated into the core model an additional negative feedback loop that acts exclusively at the transcriptional level (Fig 3A). As shown in Fig 3B, this model extension indeed improved the agreement between the mathematical model trajectories and the SOCS1 protein data (compare Fig 3B to S1B Fig) as well as the mRNA data for the selected IFNα-induced antiviral genes (compare Fig 3C to S1C Fig). Statistical analysis based on the likelihood ratio test (S2A Fig) and the Akaike information criterion (S2B Fig) confirmed that the core model with the additional intracellular feedback was significantly superior to the core model (S2C Fig).

**Fig 3:**
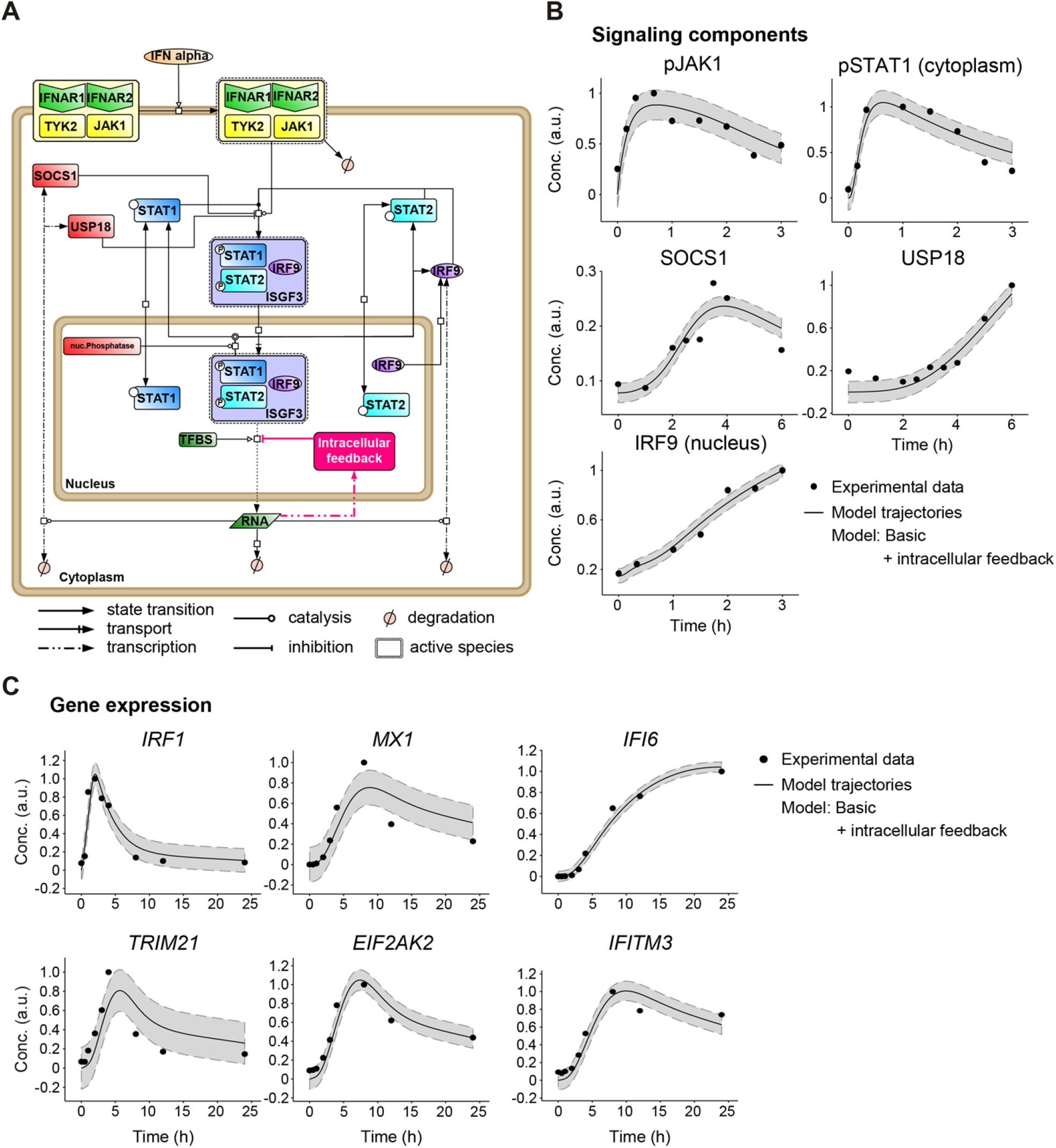
Core mathematical model with an additional intracellular feedback of IFNα-induced JAK/STAT signaling and gene expression. (**A**) Schematic representation of the core model with an additional intracellular feedback according to Systems Biology Graphical Notation. TFBS: transcription factor-binding site. (**B-C**) Trajectories of the core model with an additional intracellular feedback are shown together with the dynamic behavior of the core components of the JAK/STAT signaling pathway measured by quantitative immunoblotting (**B**) and to the expression of IFNα-induced genes examined by qRT-PCR (**C**) after stimulation of Huh7.5 cells with 500 U/ml IFNα. Filled circles: experimental data; line: model trajectories, shades: estimated error; a.u. arbitrary units.

To identify the nature of this negative intracellular factor, we performed a transcription factor binding site (TFBS) analysis using the HOMER motive discovery approach [21]. The analysis revealed six significantly enriched transcription factor binding motifs in the genes analyzed in addition to ISRE (Fig 4A), i.e. the motifs corresponding to IRF1, IRF2, IRF4, PU.1 and STAT5. Because IRF1 is a positive regulator of antiviral genes [22], this factor was excluded. IRF2 exhibits structural similarity to IRF1 [23] but possesses a repression domain and functions as a transcriptional repressor that antagonizes IRF1-induced transcriptional activation [24]. Although IRF2 and IRF4 are structurally similar, the repressive function of IRF4 was reported to be different from that of IRF2. IRF4 possesses an autoinhibition domain of DNA binding at the carboxy-terminal region that can mask the DNA-binding domain of IRF4. PU.1, as part of the Ets-transcription factor family, forms dimers with IRF4 [25]. The presence of different proteins with similar molecular functions suggests a complex network of negative regulation of IFN-induced antiviral genes and the absence of one of these factors might be compensated by the others. To quantify the impact of the identified transcription factors, we performed siRNA knock-down experiments. Expression of IRF2, IRF4 and IRF8 was downregulated by siRNA in all possible combinations and the expression levels of the selected antiviral genes were analyzed after 24 hours (S2D Fig). Interestingly, almost all combinations that included the downregulation of IRF2 positively affected gene expression. To further analyze the characteristics of the putative negative regulator of transcription, model predictions of the expression dynamics of this intracellular factor (Fig 4B) were compared with the experimentally measured mRNA expression of the selected IRFs. Only the profile of the expression kinetics of IRF2 were similar to the dynamics predicted by the model for the expression of the negative regulator (Fig 4C). Therefore, we treated Huh7.5 cells with IFNα in combination with non-targeting siRNA or siRNA directed against IRF2 and measured the expression profiles of the selected antiviral genes in a time-resolved manner. As shown in Fig 4D, knock-down of IRF2 (S2E Fig) significantly enhanced the expression of all antiviral genes monitored. These results confirmed IRF2 as an important transcriptional repressor negatively regulating IFNα-induced antiviral expression.

**Fig 4.**
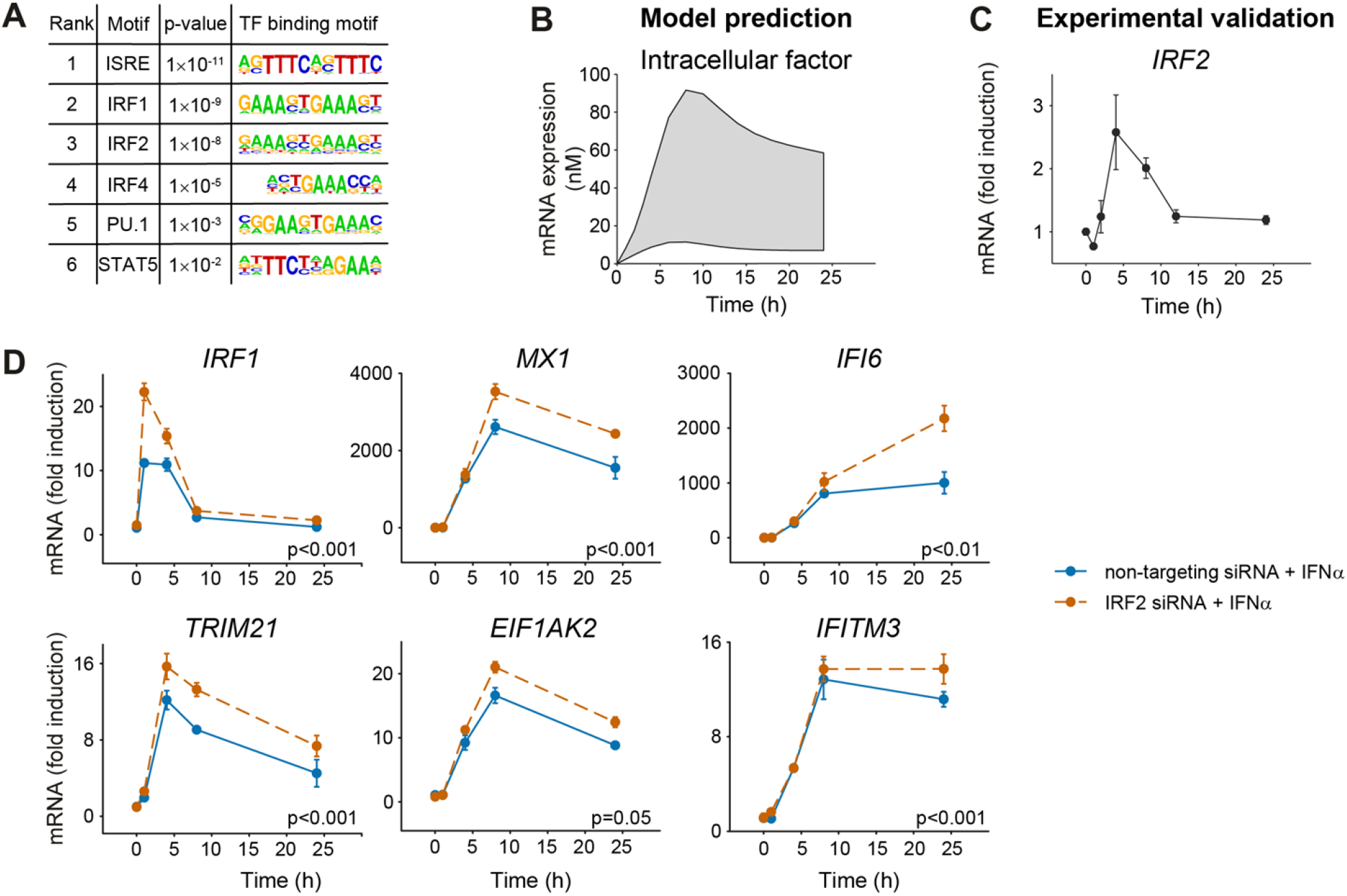
The expression profiles of the selected IFNα-stimulated genes are negatively influenced by the intracellular factor IRF2. (**A**) Transcription factor binding site analysis by the HOMER motifs software revealed six significantly regulated transcription factor binding motifs. The six most significantly enriched motifs according to the p-value and their sequence motifs are shown. (**B**) Model prediction of the mRNA expression profile of the negative regulatory intracellular factor. Shading represents the uncertainty of the prediction. (**C**) Expression profile of *IRF2* mRNA after treatment with 500 U/ml IFNα was detected by qRT-PCR. (**D**) Upregulation of gene expression by decreased IRF2 expression. Huh7.5 cells were incubated with 50 nM siRNA directed against IRF2 (orange) or non-targeting control (blue) for 24 hours, and then treated with 500 U/ml IFNα. The cells were lysed at the indicated time points and total RNA was extracted and analyzed by qRT-PCR. The error bars represent SD of biological triplicates. Significance was tested by 2-way ANOVA.

### IL1β amplifies the IFNα-induced gene response

The observation that knock-down of a negative regulator resulted in enhanced expression of early, intermediate and late IFNα-induced antiviral genes suggested that strategies could be designed to strengthen the induction of an antiviral response. Since knock-down or inhibition of an intracellular factor is difficult to achieve *in vivo*, we tested whether a similar amplified expression of IFNα-induced antiviral genes could also be mimicked by the addition of an extracellular factor. As it has been previously reported that cross-talk between IFNα and inflammatory cytokines may occur [26], we focused our analysis on inflammatory cytokines that are known to act in the liver: interleukin 6 (IL6), IL8 and IL1β. To experimentally test these cytokines, we performed co-stimulation experiments with each cytokine and IFNα and quantified the expression of the selected IFNα-induced antiviral genes in Huh7.5 cells. Co-stimulation with IL8 had no effect on the dynamics of IFNα-induced gene expression (S3A Fig), whereas treatment with IFNα and IL6 resulted in a small increase in the expression of the early gene *IRF1* (Fig 5A). Strikingly, co-stimulation with IFNα and IL1 β resulted in markedly enhanced expression of all antiviral genes examined (Fig 5B). Stimulation of Huh7.5 cells with IL1β alone resulted only in a minor increase in the expression of *IRF1* and did not elicit the expression of the other selected antiviral genes. The enhanced expression dynamics in response to co-treatment with IFNα and IL1β mimicked the effect on the expression dynamics of early, intermediate and late IFNa-induced antiviral genes observed upon knockdown of IRF2 and was even further elevated for the early antiviral gene *IRF1* and the late antiviral gene *IFITM3*. These results suggested that IL1β indeed can act as a strong amplifier of IFNα-induced expression of antiviral genes.

**Fig 5.**
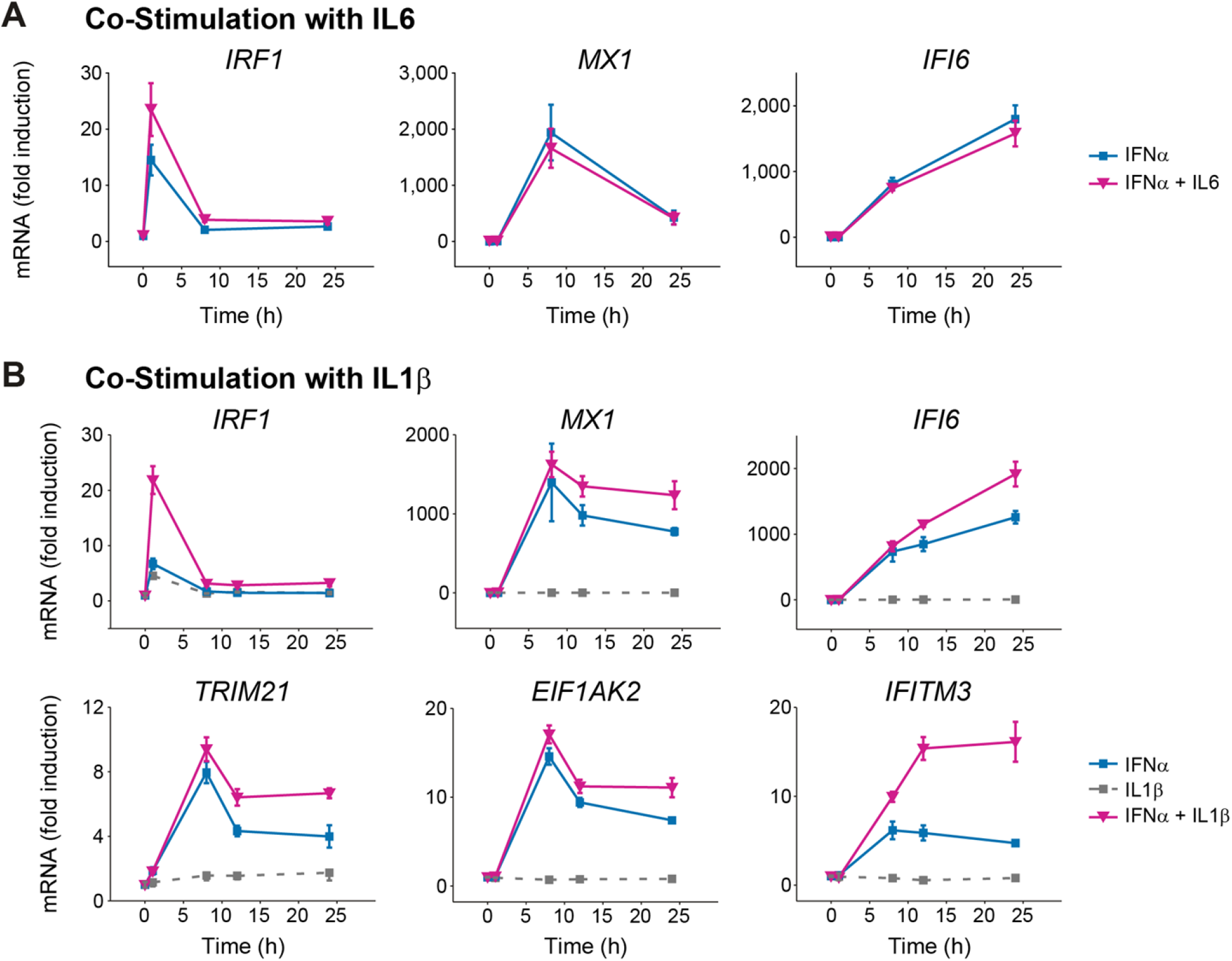
Enhanced IFNα-induced gene expression after co-stimulation with IFNα and IL1β. (A) Co-stimulation with IFNα and IL6. Huh7.5 cells were growth factor depleted followed by single treatment with 500 U/ml IFNα alone or in combination with 5 ng/ml IL6. At indicated time points RNA was extracted and analyzed using qRT-PCR. Error bars represent SD of biological triplicates. (**B**) IFNα-induced gene expression after co-treatment with IFNα and IL1β. Huh7.5 cells were treated with 500 U/ml IFNα, were stimulated with 500 U/ml IFNα alone or were co-treated with 500 U/ml IFNα and 10 ng/ml IL1β. RNA was extracted at the indicated time points and analyzed by qRT-PCR. Error bars represent SD of biological triplicates.

### IL1β-mediated STAT3 activation enhances the expression of IFNα-induced genes

It was previously reported that IL1β stimulation activates the NFκB-IκBα and the p38 signaling pathways [11]. To analyze which pathway mediated the enhancing effect of IL1β onto IFNα-induced expression of antiviral genes, Huh7.5 cells were treated with IFNα, IL1β or with a combination thereof. The dynamics of key signaling proteins in response to IFNα stimulation for up to 24 hours was analyzed by quantitative immunoblotting and for each component the area under the activation curve was calculated (Fig 6A, S3B Fig). These results showed that the phosphorylation of STAT1 was strongly induced by IFNα, but not by IL1β. However, co-treatment with IFNα and IL1β resulted in a stronger and prolonged STAT1 phosphorylation. Single IL1β treatment or co-stimulation with IFNα induced the activation of the p38 pathway and p65 of the NFκB pathway to a similar extent. Strikingly, phosphorylation of STAT3 was detected after stimulation with IL1β alone as well as after IL1β and IFNα co-treatment, whereas IFNα alone only resulted in a weak activation of STAT3. The comparison of the area under the curve of STAT3 phosphorylation showed that STAT3 phosphorylation was significantly increased in the IL1β and IFNα co-treated samples. To assess whether the increased phosphorylation of STAT3 correlated with nuclear accumulation of STAT3 in particular at late time points, we performed live cell imaging experiments with primary hepatocytes from an *mKate-Stat3* knock-in mouse strain expressing a fluorescently tagged STAT3 [27] (Fig 6B). Compared to the treatment with IL6 that resulted in an instantaneous nuclear translocation of STAT3 (S3C Fig), nuclear STAT3 was detectable at lower levels and at later time points in response to IL1β stimulation. However, it was markedly elevated upon co-treatment with IFNα and IL1β at later time points, in particular 24 hours post treatment (Fig 6B). Therefore, the sustained STAT3 phosphorylation profiles and the nuclear accumulation of STAT3 observed upon co-treatment with IFNα and IL1β matched the co-stimulatory effect of IL1β and IFNα on the expression of the selected IFNα-induced antiviral genes.

**Fig 6.**
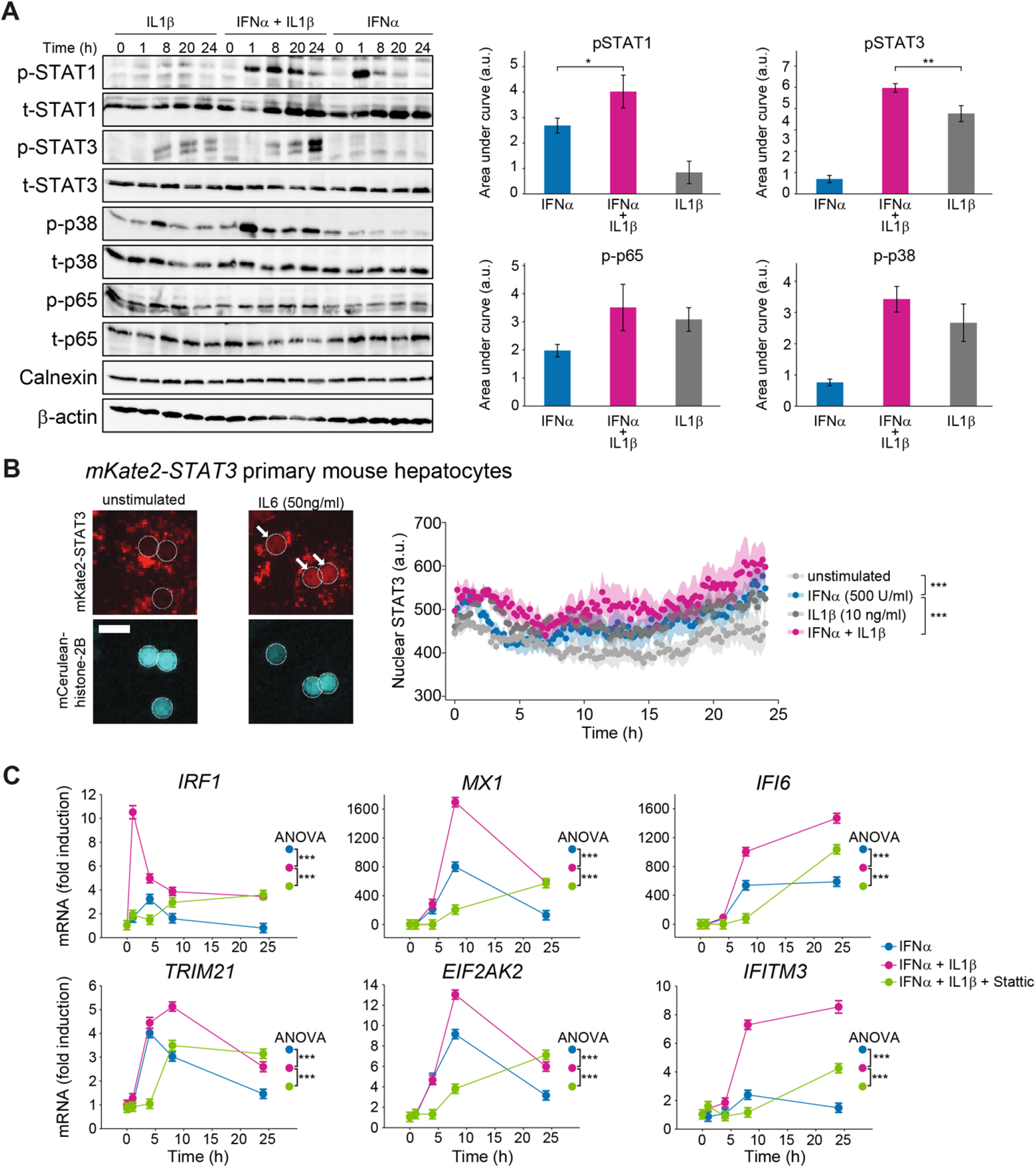
Co-stimulation of IFNα and IL1β results in phosphorylation of STAT3. (**A**) Huh7.5 cells were single or co-stimulated with 500 U/ml IFNα and 10 ng/ml IL1β. Cells were lysed at indicated time points and analyzed using quantitative immunoblotting. Error bars represent SEM of three biological replicates. (**B**) Primary mouse hepatocytes from *mKate2-Stat3* knock-in mice were growth factor depleted overnight and stimulated with IL6 or left untreated. Representative images of cells expressing a mCerulean-histone-2B nuclear marker are depicted. The dotted line indicates the outline of the nuclei and white arrows indicate nuclear STAT3. The right panel indicates the quantification of the nuclear mKate2-STAT3 intensity of primary mouse hepatocytes stimulated with IL1β, IFNα, co-stimulated or left unstimulated. The quantification is based on 20 cells per condition; shading indicates SEM; scale bar=25 μm. Significance was tested by two-way ANOVA, ***, p<0.001. (**C**) Huh7.5 cells were pre-treated for 30 minutes with 10 μM STAT3 inhibitor Stattic followed by 500 U/ml IFNα in combination with 10 ng/ml IL1β. mRNA was extracted at indicated time points and analyzed using qRT-PCR. Error bars represent SD of biological triplicates.

To ascertain that STAT3 activation contributes to the enhanced expression of the selected IFNα-induced antiviral genes, single or co-stimulated Huh 7.5 cells were either left untreated or were co-treated with a STAT3 inhibitory compound (Stattic) [28]. Treatment of Huh7.5 cells with 10 μM Stattic for up to 24 hours had no significant impact on their viability (S3D Fig). With this dose of Stattic, the induction of STAT3 phosphorylation by co-stimulation with IFNα and IL1β was reduced for the entire observation time (S3E Fig). Analyzing gene expression, we noticed that at the early time points the expression of all selected genes induced by IFNα and IL1β co-stimulation was reduced by treatment with Stattic (Fig 6C). At 24 hours after IFNα and IL1β co-stimulation, expression of both early and intermediate IFNα-induced antiviral genes was comparable for Stattic-treated and untreated samples. However, the late IFNα-induced genes, *IFI6* and *IFITM3*, showed a strong decrease in their expression upon Stattic treatment during the entire observation time (Fig 6C). Overall, application of the STAT3 inhibitor Stattic had a significant effect on the expression of all analyzed IFNα-induced antiviral genes. These results indicated that co-stimulation of cells with IFNα and IL1β enhanced the activation of STAT3, thus mediating the amplified expression kinetics of IFNα-induced antiviral genes.

### IL1β enhances IFNα-induced gene expression in primary human hepatocytes and viral clearance

To assess whether the IL1β-induced amplification of IFNα-induced expression of antiviral genes was conserved in primary human hepatocytes and relevant for eliciting an antiviral response, we first examined the impact of IL1β on the dynamics of IFNα-induced antiviral genes in these cells. As shown in Fig 7A, consistent with our observations in Huh7.5 cells, co-stimulation of primary human hepatocytes with IFNα and IL1β increased the expression especially of the early IFNα-induced gene IRF1 and the late gene IFI6. These results underscored the importance of our findings also in the context of primary human hepatocytes.

**Fig 7.**
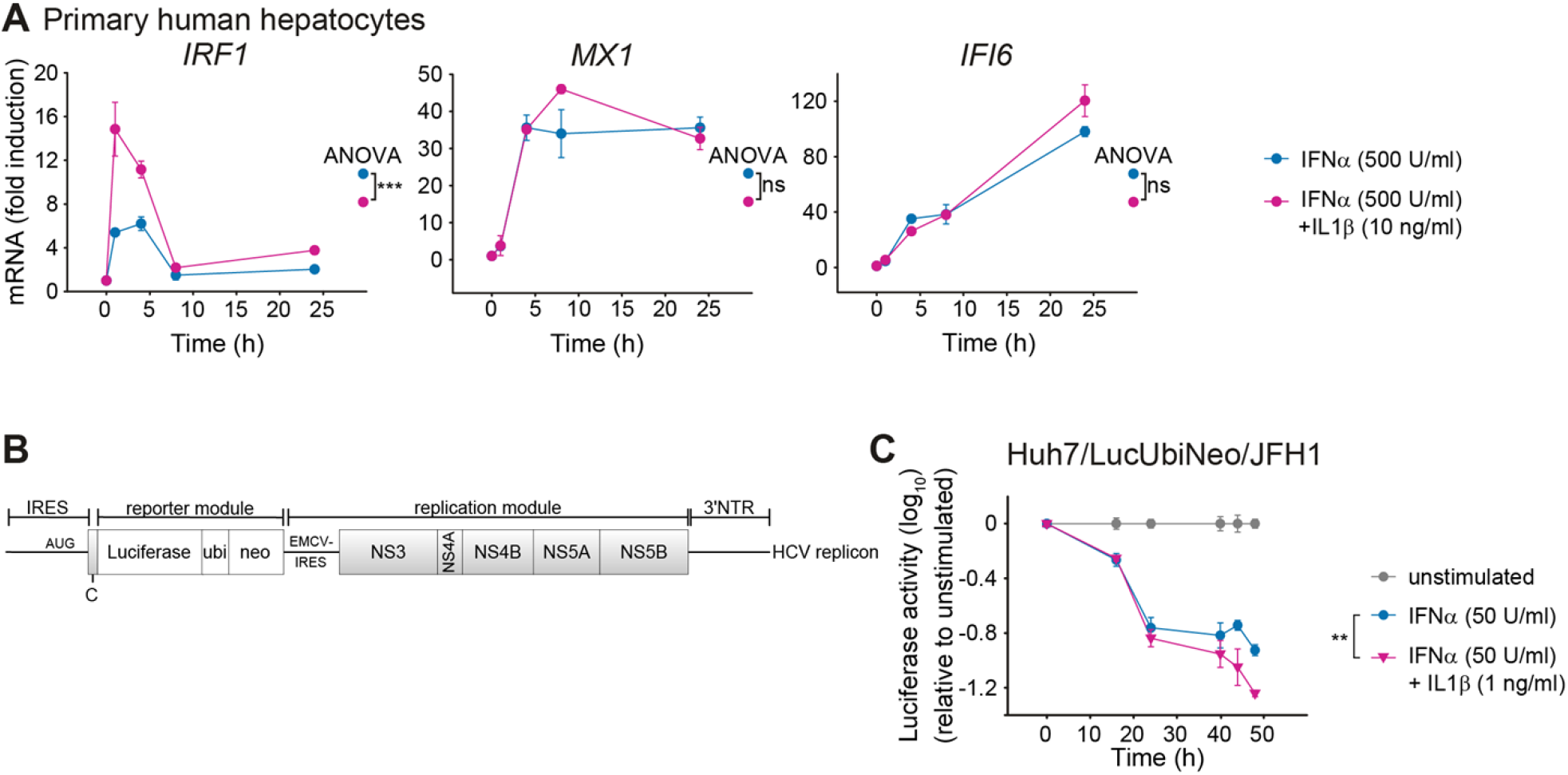
Co-stimulation with IFNα and IL1β enhances IFNα-induced gene expression in primary human hepatocytes and viral clearance of HCV in a replicon cell line. (**A**) Effect of co-stimulation with IFNα and IL1β on mRNA expression of IFNα-induced genes in primary human hepatocytes. Cells were stimulated with 500 U/ml IFNα alone or co-treated with 500 U/ml IFNα and 10 ng/ml IL1β. RNA was extracted at the indicated time points and analyzed by qRT-PCR. Samples were analyzed from three different donors and error bars represent SD. (**B**) Scheme of the bicistronic subgenomic HCV reporter RNA (replicon). IRES, internal ribosomal entry site; NTR, non-translated region. (**C**) Enhanced suppression of HCV replication in cells co-stimulated with IFNα and IL1β. Huh7/HCV/Luc replicon cells were stimulated with 50 U/ml IFNα alone or were co-treated with 50 U/ml IFNα and 1 ng/ml IL1β. The values are relative to the unstimulated control. Error bars represent the SEM of six technical replicates.

Next, we examined whether the increased expression of IFNα-induced antiviral genes in response to co-treatment with IFNα and IL1β resulted in enhanced viral clearance. For these studies we utilized a cell line containing a persistently replicating HCV reporter replicon (Huh7/LucUbiNeo/JFH1) (Fig 7B). In this cell line, luciferase activity correlates linearly with viral replication [29]. Treatment of the replicon cells with 500 U/ml IFNα – an IFNα dose that was employed in the experiment examining activation of signaling pathways or expression of antiviral genes – resulted in a very rapid inhibition of HCV replication. At this dose, a detectable but not major difference between treatment with IFNα alone and the co-stimulation with IFNα and IL1β was observed (S4A Fig). To increase the resolution of the assay and taking into account the high IFNα-sensitivity of HCV, the applied IFNα and IL1β concentrations were reduced 10-fold. In this setting, co-stimulation with IFNα and IL1β resulted in a stronger reduction in luciferase activity than IFNα alone, especially at later time points (>24 hours) (Fig 7C and S4B Fig). In conclusion, IL1β enhanced the antiviral effect of IFNα treatment and reduced HCV replication.

### IL1β-mediated enhanced expression of IFNα-induced genes requires the IL1β receptor

To confirm the specificity of the observed augmentation of the IFNα response by IL1β, primary mouse hepatocytes were isolated from wildtype and from mice lacking the IL1 receptor (IL1R1^−/−^ mice) [30]. Expression analysis of the selected IFNα-induced genes upon treatment with 500 U/ml murine IFNα or 10 ng/ml murine IL1β confirmed that treatment with IL1β alone did not induce expression of the selected IFNα-induced genes, whereas IFNα stimulation significantly upregulated their expression (Fig 8A). Co-stimulation with IFNα and IL1β synergistically increased the expression of the selected IFNα-induced antiviral genes. These experiments revealed that mRNA expression profiles of the selected IFNα-induced antiviral genes in primary mouse hepatocytes are comparable to those in Huh7.5 cells. Of note, while IFNα-induced expression of the selected IFNα-induced antiviral genes in hepatocytes from IL1R1^−/−^ mice lacking IL1β signaling was comparable to wildtype cells, IL1R1^−/−^ cells did not show a synergistic enhancement of IFNα−induced gene expression upon co-stimulation with IL1β. These results confirmed that the co-stimulatory effect of IL1β on the IFNα-induced antiviral response is mediated by the IL1R1 and that the underlying mechanism is conserved in mouse and human.

**Fig 8.**
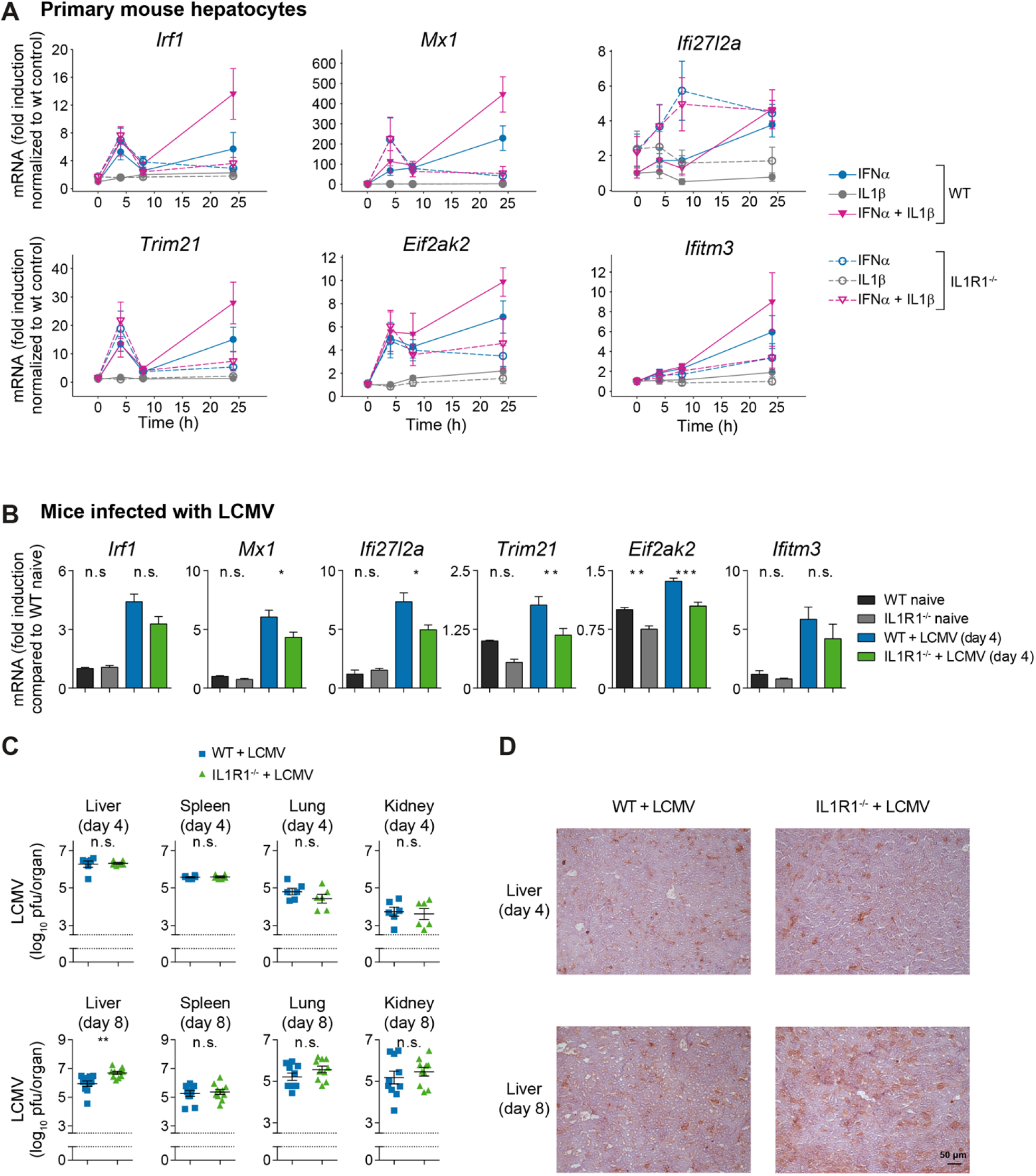
IFNα-induced antiviral response is reduced and virus replication is enhanced in IL1R1^−/−^ mice. (**A**) Expression of the selected antiviral genes in primary mouse hepatocytes from wild-type (WT) or IL1R1 knock-out (IL1R1^−/−^) mice upon stimulation with 500 U/ml murine IFNα2, 10 ng/ml murine IL1β or co-treatment. RNA was extracted at the indicated time points and analyzed by qRT-PCR. Error bars represent SD of four biological replicates; a.u.: arbitrary units. (**B**) Wild-type (wt) or IL1R1 knock-out (IL1R1^−/−^) CL57BL/6 mice were infected with 2×10^6^ pfu of LCMV strain WE. Prior to and four days post infection, livers were isolated and the selected antiviral genes were measured by qRT-PCR. Differences between WT and IL1R1^−/−^ livers were tested by one-way analysis of variance. ***, p<0.001; **, p<0.01; *, p<0.05; n.s., not significant; n=6. (**C**) Wild-type (wt) or IL1R1 knock-out (IL1R1^−/−^) CL57BL/6 mice were infected with 2.10^6^ pfu of LCMV WE. Four and eight days post infection, livers, spleens, lungs and kidneys were isolated and viral load was quantified. Titer differences between WT and IL1R1^−/−^ organs were tested by two-sided t-tests. **, p<0.01; n.s., not significant; n=6-10. (**D**) Wildtype (wt) or IL1R1 knock-out (IL1R1^−/−^) CL57BL/6 mice were infected with 2.10^6^ pfu of LCMV WE. Four and eight days past infection, livers were isolated and viral proteins (LCMV-NP) were stained; n=6-7; scale bar=50 μm.

### Viral infection is enhanced in IL1R1^−/−^ mice

To demonstrate the *in vivo* relevance of our findings, wildtype and IL1R1^−/−^ mice [30] were infected with 2×10^6^ pfu of LCMV stain WE. Prior to and four days post infection, the expression of the selected IFNα-induced genes in the liver of the animals was determined by qRT-PCR. In line with our hypothesis, the mRNA concentrations of the IFNα-induced genes *Mx1*, *Ifi27l2a*, *Trim21* and *Eif2ak2* were significantly reduced in infected IL1R1^−/−^ mice compared to wildtype mice (Fig 8B). This was not due to differences in viral load, as comparable virus amounts were detected four days post infection (Fig 8C). However, in line with the reduced antiviral response in the liver, we observed a significant increase in LCMV titers in the liver of IL1R1^−/−^ mice as compared to wildtype controls eight days post infection (Fig 8C). Consistently, immunohistochemical evaluation of liver tissue revealed that LCMV nucleoprotein (NP) was more abundant in hepatocytes of IL1R1 deficient mice than in wild type counterparts (Fig 8D). Notably, antiviral T-cell immunity was also reduced eight days post infection following LCMV infection in IL1R1^−/−^ compared to control animals (S5A-C Fig). In conclusion, the *in vivo* experiments confirmed the importance of the IL1β induced signal transduction mediated by the IL1 receptor for enhancing the IFN-induced response.

## Discussion

We observed that the temporal expression profiles of IFNα-induced genes can be classified into three different groups based on the time point of maximal activation: early, intermediate and late. By mathematical modeling based on time-resolved experimental data, our studies revealed that mRNA stability and expression of IRF2 as a negative regulator of transcription critically determine the expression profiles of IFNα-induced genes. Strikingly, we observed that IL1β can mimic the impact of IRF2 knockdown and significantly boost IFNα-induced responses.

It has previously been reported that TNF stimulation of mouse fibroblasts for twelve hours resulted in early, intermediate and late gene expression clusters and that these clusters differ in mRNA stability [31]. Consistent with these observations, we demonstrated that the mRNA half-lives of the IFNα-induced antiviral genes indeed differ substantially among the three groups and correlate with their peak of expression.

Positive and negative feedback mechanisms establish a balanced regulatory network of type I IFN-induced signaling [1] and the combination of transcriptional activators and repressors is critical for the expression of specific genes and viral clearance. Consistent with previous results [24], we observed that the transcription factor IRF2 is induced by IFNα. In addition, we demonstrated that the downregulation of IRF2 by siRNA enhances antiviral gene expression, which is in agreement with an elevated IFN-induced gene expression in IRF2-deficient mice [32]. Furthermore, it has been demonstrated that IRF2 knock-down results in the upregulation of IFN-induced genes in the bone marrow [33]. Virus-induced IFNβ expression is substantially higher in IRF2-deficient mice than in wild-type mice [34], and HCV-infected patients exhibit increased expression of IRF2 [35]. In line with these observations, we showed that IRF2 negatively regulates the expression of IFNα-induced genes and represents an important feedback mechanism dampening the type I IFN response.

Additionally, we provided evidence that co-stimulation with IL1β enhances the expression of IFNα-induced genes. In agreement with this observation, it was previously reported that IFNα and IL1β co-stimulation in Huh7.5 cells increased the phosphorylation of STAT1 and resulted in an increased expression of two antiviral proteins, PKR (encoded by *EIF2AK2*) and OAS, compared to treatment with IFNα alone [18]. In our study, co-stimulation with IFNα and IL1β rather shifted the peak of STAT1 phosphorylation to later time points.

We further showed that IL1β stimulation strikingly induced the phosphorylation of STAT3 at time points later than 6 hours. The IL1β-induced activation profile of STAT3 was remarkably different from the IL6-induced STAT3 phosphorylation that peaks at one hour after stimulation. At present, there are only very few reports on STAT3 activation by IL1β. For example, IL1β-induced phosphorylation of STAT3 was reported in myocytes [36], in mesangial cells [37] and in HepG2 cells with a weak increase of phosphorylation eight hours after stimulation [12]. In our study, inhibition of STAT3 by the treatment with the inhibitor Stattic reduced the phosphorylation of IL1β-induced STAT3 activation and the expression of antiviral genes after IFNα and IL1β co-stimulation. Likewise, in RAW 264.7 cells, a reduction of LPS-induced STAT3 activation and target gene expression was observed upon treatment with the inhibitor Stattic [38]. In conclusion, this is to our knowledge the first report indicating that IL1β stimulation triggers prolonged STAT3 phosphorylation and nuclear translocation.

Although IL1β on its own did not affect the expression of IFNα-induced genes in the observed time frame of 24 h, co-treatment with IFNα elevated their expression and enhanced for example the antiviral state as inferred from the increased inhibition of HCV replication. Moreover, in LCMV infection *in vivo,* viral titers were increased in IL1R-knock-out mice, showing that IL1β signaling through this receptor contributes to viral clearance. Clinical data demonstrated that levels of pro-inflammatory cytokines including IL1β, IL4 and IL6 are elevated in the sera of patients with HCV infection [39]. However, the role of IL1β in hepatitis virus-infected individuals and the impact on viral clearance are controversially discussed. On the one hand, it was reported that IL1β concentrations are within the normal range during IFNα treatment of HCV patients [40] and decrease in chronically infected patients [41]. On the other hand, Daniels *et al.* demonstrated that the increased production of IL1β by peripheral blood mononuclear cells during IFNα treatment contributes to the inhibition of hepatitis B virus replication and promotes viral clearance [42]. Similarly, Zhu *et al.* reported that IL1β inhibits HCV replication in a hepatoma-derived replicon cell line [43].

In conclusion, we demonstrate that IL1β boosts the expression of IFNα-induced antiviral genes, and in vivo particularly those with an intermediate and a late expression profile. IL1β thereby could strengthen the efficacy of therapeutically applied IFNα in particular in the liver and this knowledge might help to improve IFN-based strategies for the treatment of viral infections.

## Materials and Methods

### Cell Culture

Huh7.5 cells were kindly provided by Charles M. Rice (The Rockefeller University, NY, RRID:CVCL_7927) and primary human hepatocytes (PHH) were kindly provided by Georg Damm (Charité Berlin). Murine hepatocytes were isolated from wildtype or from IL1R1^−/−^ CL57BL/6 mice as previously described [44].

All cells were cultivated at 37°C and 5 % CO_2_ incubation and 95 % relative humidity. Informed consent of the patients for the use of tissue for research purposes was obtained corresponding to the ethical guidelines of the Charité-Universitätsmedizin Berlin. The Huh7.5 cell line was authenticated using Multiplex Cell Authentication and the purity of cell line was validated using the Multiplex Cell Contamination Test by Multiplexion (Heidelberg, Germany) as described recently [45, 46].

### Cells stimulation for protein and mRNA measurements

One day before time-course experiments, 1.7.10^6^ Huh7.5 cells or 2.10^6^ PHH were seeded intoa 6 cm-diameter dishes or 5.5.10^5^ cells per well of 6-well plates in culture medium. Huh7.5 were cultured in Dulbeccos’s Modified Eagle Medium (DMEM, Invitrogen) supplemented with 10% fetal calf serum (FCS) (Gibco) and 1% P/S (Invitrogen). PHHs were cultivated in Williams medium E (Biochrom) supplemented with 10% FCS (Gibco), 100 nM dexamethasone, 10 μg/ml insulin, 2 mM L-Glutamin (Gibco) and 1% Penicillin-Streptomycin (P/S) (Invitrogen). Prior to stimulation, cells were washed three times with PBS and cultivated in serum free medium for three hours. Stimulation of cells was performed by adding the stimulation factor directly into serum free medium. To stop stimulation, dishes were placed on ice, medium was aspirated and cells were lysed either with Nonidet P-40 lysis buffer (1% NP40, 150 mM NaCl, 20 mM Tris pH 7.4, 10 mM NaF, 1 mM EDTA pH 8.0, 1 mM ZnCl_2_ pH 4, 1 mM MgCl_2_, 1 mM Na_3_VO_4_, 10% Glycerol and freshly added 2 μg/ml aprotinin and 200 μg/ml AEBSF) or Nonidet P-40 cytoplasmic lysis buffer (0,4% NP40, 10 mM HEPES pH 7.9, 10 mM KCl, 0.1 mM EDTA, 0.1 mM EGTA and freshly added 2 μg/ml aprotinin, 200 μg/ml AEBSF, 1 mM DTT, 1 mM NaF and 0.1 mM Na_3_VO_4_) and nuclear lysis buffer (20 mM HEPES pH 7.9, 25% glycerin, 400 mM NaCl, 1 mM EDTA, 1 mM EGTA and freshly added 2 μg/ml aprotinin, 200 μg/ml AEBSF, 1 mM DTT, 1 mM NaF and 0.1 mM Na_3_VO_4_) for cell fractionation. To measure the viability of cells upon Stattic treatment, CellTiter-Blue Viability Assays (Promega) were performed according to the manufacturer’s instructions. Incubation with the dye for 60 min was followed by measurement of the fluorescence with the infinite F200 pro Reader (Tecan).

### RNA analysis

Cells were seeded, growth factor depleted and stimulated with IFNα (PBL, 11350-1). Total RNA was isolated from three independent dishes per time point by passing the lysate through a QIAshredder (Qiagen) for homogenization, followed by RNA extraction using the RNeasy Plus Mini Kit (Qiagen) according to manufacturer’s protocol. For cDNA generation, 1 μg of total RNA was used and transcribed with the High-Capacity cDNA Reverse Transcription Kit (Applied Biosystems) according to manufacturer’s instructions. Quantitative real-time PCR (qRT-PCR) was performed using the hydrolysis-based Universal Probe Library (UPL) platform (Roche Diagnostics) in combination with the Light Cycler 480 (Roche Diagnostics). Primers were generated using the automated Assay Design Center based on species and accession number (www.lifescience.roche.com) (see S2 Table). Crossing point (CP) values were calculated using the second derivative maximum method of the Light Cycler 480 software (Roche Diagnostics). An internal dilution series of template cDNA (stimulated for 1 hour with 500 U/ml IFNα) was measured with every gene analyzed for PCR efficiency correction and served as standard curve for calculation of relative concentrations. Relative concentrations were normalized to HPRT.

### Quantitative immunoblotting

For Immunoprecipitation (IP), the target-specific antibody was added to the cellular lysates together with 25 μl of Protein A or G sepharose (GE Healthcare) depending on the species of target antibody and the mixture was incubated overnight rotating at 4°C. For anti-JAK1 (Upstate Millipore, 06-272, RRID:AB_310087), anti-Tyk2 (Upstate Millipore, 06-638, RRID:AB_310197) and anti-STAT1 (Upstate Millipore, 06-501, RRID:AB_310145) IP, Protein A sepharose was used. Protein G sepharose was used for anti-SOCS1 (Millipore, 04-002, RRID:AB_612104) IP. Protein concentration of cellular lysates was determined using the BCA Assay kit (Pierce/Thermo Scientific) according to the manufacturer’s instructions. Proteins were separated by denaturing 10% or 15% SDS-PAGE. Sample loading was randomized to avoid systematic errors [47]. The proteins were transferred to PVDF (STATs, IRF9, USP18) or nitrocellulose membranes (JAK1, TYK2). Membranes were stained with 0.1% Ponceau Red (Sigma-Aldrich). To detect tyrosine phosphorylation of immunoprecipitated JAK1 and TYK2, the anti-phosphotyrosine monoclonal antibody 4G10 (Upstate Biotechnology, 05-321, RRID:AB_309678) was used. Phosphorylation specific and total antibodies:

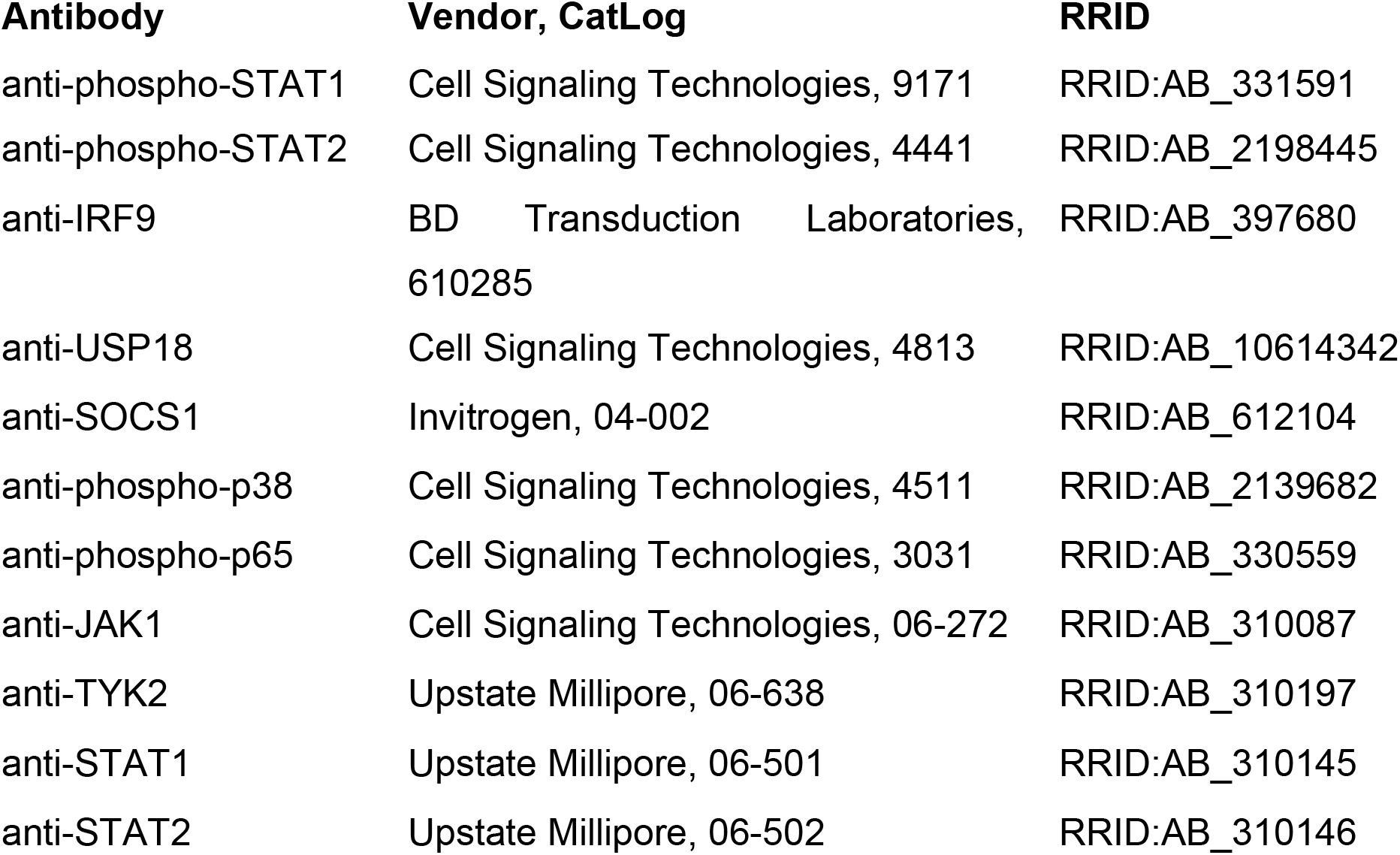
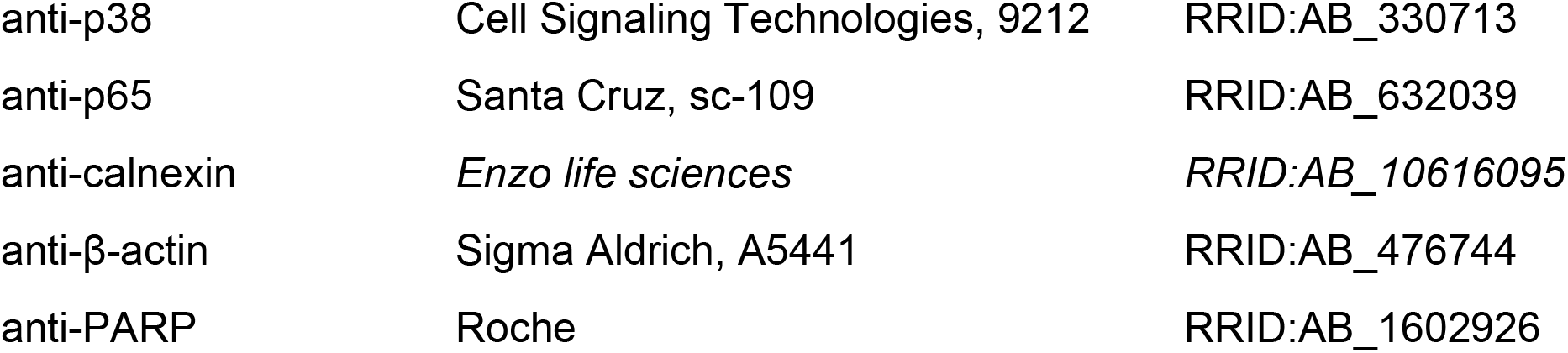

For detection of additional proteins on the same membrane, membranes were incubated with β-mercaptoethanol and SDS. For normalization, antibodies against calnexin and β-actin were used for the cytoplasmic fraction and anti-PARP was used for the nuclear fraction. Secondary horseradish peroxidase-coupled antibodies (anti-rabbit HRP, anti-goat HRP, Protein A HRP) were purchased from GE Healthcare. Immunoblots were incubated with ECL or ECL advance substrate (GE Healthcare) and signals were detected with a CCD camera (ImageQuant LAS 4000 biomolecular imager (GE Healthcare)). Immunoblot data was quantified using ImageQuant TL version 7.0 software (GE Healthcare). Quantitative immunoblot data were processed using GelInspector software [47]. Data normalization was performed by using either the recombinant calibrator proteins GST-JAK1DN or GST-Tyk2DC for JAK1 and TYK2, respectively, or housekeeping proteins: β-actin for IRF1, IRF9, USP18, p38 and p65 or calnexin and PARP for STAT1 and STAT2 in the cytoplasm and nucleus, respectively. For smoothing splines to the data, Matlab’s csaps-splines with a smoothing parameter of 0.8 were used.

### siRNA transfection

For siRNA transfection, 2.25×10^5^ Huh7.5 cells were seeded in 6-well plates 24 hours prior to transfection. The next day, cells were washed three times with PBS and cultivated in P/S free DMEM supplemented with 10% FCS before the transfection with 50 nM siRNA (Dharmacon) (IRF2: L-011705-02-0005; non-targeting siRNA: D-001810-10-20). Transfection was performed by incubation of siRNA with Optimem Medium (Gibco, Life Technologies) and Lipofectamin RNAiMAX (Invitrogen) for 20 minutes at RT and adding the mixture dropwise to cells. For efficient uptake, cells were incubated with siRNA transfection mixture for 24 hours. Subsequently, the medium was changed and time course experiments were performed.

### Live-cell imaging

Primary hepatocytes (15,000 cells per well, 96-well plate format) derived from *mKate2-STAT3* heterozygous knock-in mice [27] were transduced with adeno-associated viruses encoding mCerulean-labeled histone-2B during adhesion. Cells were cultivated as described above, stimulated with ligand, and imaged using a Nikon Eclipse Ti Fluorescence microscope in combination with NIS-Elements software. Temperature (37°C), CO_2_ (5%) and humidity were held constant through an incubation chamber enclosing the microscope. Three channels were acquired for each position: bright-field channel, STAT3 channel (mKate2), and nuclear channel (CFP). Image analysis was performed using Fiji software, and data were processed using R software.

### Luciferase assay

Luciferase activity was measured as read out for HCV replication. 30,000 cells of the replicon cell line Huh7/LucUbiNeo/JFH1 [48] were seeded in a 24-well plate two days prior to the stimulation. Cells were growth factor depleted for 3 hours followed by IFNα treatment. At different time points cells were washed once with PBS and lysed with 100 μl luciferase lysis buffer (1% Triton X-100, 25 mM glycil-glycin (pH 7.8), 15 mM MgSO_4_, 4 mM EGTA, 10% Gylcerol) directly in the well. Plates were stored at −80°C until measurement. Luciferase was measured applying 400 μl luciferase assay buffer (15 mM K_3_PO_4_ (pH7.8), 25 mM glycil-glycin (pH 7.8), 15 mM MgSO_4_, 4 mM EGTA) with freshly added 1 mM DTT, 2 mM ATP and 1mM D-Luciferin. Luciferase activity was measured using Mitras^2^ multimode reader LB942 (Berthold).

### Cultivation of primary mouse hepatocytes

Cells were seeded with a density of 3.5×10^5^ cells per cavity in a collagen-coated 6-well-plate. For the experiments cells were cultivated under FCS-free conditions in DMEM/Ham’s F-12 (Biochrom) supplemented with 2 mM glutamine and 100 U/ml penicillin/0.1 mg/ml streptomycin (Cytogen). Cells were stimulated with 500 U/ml of murine recombinant IFNα (PBL) with or without 10 ng/ml of murine recombinant IL1β (JenaBioscience) for the time points indicated in the respective figure.

### RNA isolation and qRT-PCR of primary mouse hepatocytes

Total cellular RNA was isolated by using the RNeasy Miniprep Kit (Qiagen) as described in the manufacturer’s instructions. 1 μg of total RNA was reverse transcribed with Quantitect Reverse Transcription Kit (Qiagen) using oligo(dT), which included DNase I digestion. cDNA was diluted 1/5, and 1.2 μl of the diluted cDNA was added as template to a final volume of 25 μl including 1x GoTaq qPCR Master Mix according to the manufacturer’s instructions (Promega, Mannheim, Germany). qRT-PCR was performed using the ViiA7 real-time PCR system (Applied Biosystems). Primers were generated using the Primer-BLAST design tool from NCBI based on the accession number of the gene of interest. All primers were purchased from Eurofins MWG Operon (Ebersberg). Specificity of rtPCR was controlled by no template and no reverse-transcriptase controls. Semiquantitative PCR results were obtained using the ΔCT method. As control gene *HPRT* was used. Threshold values were normalized to *HPRT* respectively.

RNA purification of liver tissue from LCMV infected mice for qRT-PCR analyses were performed as previously described [49]. Gene expression of *IRF1*, *MX1*, *ISG12*, *TRIM21*, *EIF2AK2*, *IFITM3*, *HPRT* was performed using kits from Applied Biosystems. For analysis, the expression levels of all target genes were normalized to *HPRT* expression (ΔCt). Gene expression values were then calculated based on the ΔΔCt method, using the naïve liver samples as a control to which all other samples were compared. Relative quantities (RQ) were determined using the equation: RQ=2^−ΔΔCt^.

### LCMV infection of wild-type or IL1R1 knock-out mice

All mice were on a C57BL/6 genetic background. IL1R1^−/−^ mice [30] were obtained from Jackson Laboratory (mouse strain 003245). All mice were maintained under specific pathogen-free conditions and experiments have been approved by the LANUV in accordance with German laws for animal protection (reference number G315). LCMV strain WE was originally obtained from F. Lehmann-Grube (Heinrich Pette Institute, Hamburg, Germany) and was propagated in L929 cells as described. Mice were infected intravenously with 2×10^6^ plaque forming units (pfu) LCMV-WE. Virus titers were measured using a plaque forming assay as described previously [49]. Briefly, organs were harvested into HBSS and homogenized using a Tissue Lyser (Qiagen). 0.8×10^6^ MC57 cells were added to previously in 10-fold dilutions titrated virus samples on 24-well plates. After 3h 1% methylcellulose containing medium was added. After 48 h plates were fixed (4% formalin), permeabilized (1% Triton X HBSS), and stained with anti-VL-4 antibody, peroxidase anti-rat antibody and PPND solved in 50 mM Na_2_HPO_4_ and 25 mM citric acid. Histological analysis was performed on snap frozen tissue as described [49]. Anti-LCMV-NP (clone: VL4) was used in combination with an alkaline phosphatase system. Tetramer production, surface and intracellular FCM staining was performed as described previously [49]. Briefly, single cell suspensions from spleen and liver tissue as well as peripheral blood lymphocytes were stained using gp33 or np396 MHC class I tetramers (gp33/H-2Db) for 15 min or gp61 MHC II tetramer for 30 min at 37°C, followed by staining with anti-CD8 (BD Biosciences) for 30 min at 4°C. For determination of their activation status, lymphocytes were stained with antibodies against surface molecules as indicated for 30 min at 4°C. For intracellular cytokine stain single suspended splenocytes or liver cells were incubated with the LCMV-specific peptides gp33, np396, or gp61. After 1 h Brefeldin A (eBiosciences) was added, followed by additional 5 h incubation at 37°C. After surface stain with anti-CD8 or anti-CD4 (eBiosciences) cells were fixed with 2% formalin and permeabilized with PBS containing 1% FCS and 0.1% Saponin and stained with anti-IFNγ (eBiosciences) for 30 min at 4°C.

### Microarray analysis

IFNα-induced gene expression data [4] was analyzed by the Robust Multi-array Average (RMA) [50] algorithm. It was applied for data processing of Affymetrix gene expression data (Human Gene ST Arrays) using the implementation in the simpleaffy R package version 2.40.0 (http://www.bioconductor.org/packages/release/bioc/html/simpleaffy.html). All subsequent analyses were performed on the log_2_-scale and the expression of the individual genes was considered relative to the measured expression of untreated cells at 0 hours. A paired t-test (treated vs. untreated) was used to assess the significance of IFNα-induced regulation at 1, 2, 3, 4, 8, 12, 24 hours. Because only three genes (ID8139776, TCEB3CL2, CFC1) were significantly downregulated, we focused on the 53 genes showing a significant upregulation (p<0.05 and average fold-change>2). The time-point of maximal regulation was considered to subdivide the upregulated genes into three classes. Genes were visualized with respect to the time-point of maximal regulation and within the groups with the same time-point according to the fold change at 1 h.

### Quantification of RNA stability

Cells were seeded, growth factor depleted and stimulated with 500 U/ml IFNα for 8 hours as described above followed by treatment with 5 μg/ml actinomycin D to inhibit transcription. Total RNA was extracted at specific time points and analyzed using qRT-PCR. RNA half-life was estimated by fitting the mRNA fold expression to an exponential decay 3-parameter function. t_1/2_: mRNA half-life.

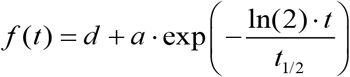

### Quantification of dose-dependency of RNA on IFNα

Cells were seeded, growth factor depleted and stimulated as described above and stimulated with increasing doses of IFNα for 4 hours. Total RNA was extracted and analyzed using qRT-PCR. A sigmoidal 4-parameter Hill function was fitted to the RNA expression.

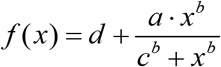

Where *d* = y-axis intercept, *a* = amplitude, *b* = slope and *c* = x-value of the point of inflection i.e. the EC_50_ dose.

### Transcription factor binding site analysis

Transcription factor binding site analysis was performed using HOMER software [21] (http://homer.salk.edu/homer/ngs/index.html). Promoter regions of analyzed genes were analyzed for known transcription factor binding site. For this, a list with identifiers of genes of interest was submitted to the software and the respective promoter regions were obtained from a software-specific database. Significant enrichment of found transcription factor binding sites were set relative to all promoter regions analyzed using hypergeometric test.

### Mathematical modeling

The presented modeling approach is based upon a previously published IFNα model [4]. For this study, the model has been extended by incorporating the genetic response of IFNα-stimulated JAK/STAT signaling. Further, the formation of the receptor complex was simplified so that the complex is activated directly by IFNα binding. In addition, ISGF3 formation and dissociation were previously incorporated as two steps. Here, the complete formation of ISGF3 was summarized in one step; STAT1, STAT2 and IRF9 bind synergistically. The final model consists of 30 species and 53 kinetic parameters. All reactions are defined as ordinary differential equations (ODEs) based on mass action kinetics in cytoplasm and nucleus. Measured concentrations (STAT1, STAT2, IRF9 and IFN) were transformed from molecules per cell to nM by using STAT1 concentration as reference. In the final version of the model, unphosphorylated STAT1 concentration was identified to be negligible. The current model is implemented into the MATLAB-based modeling framework D2D [51, 52].

### Parameter estimation

To find the optimal parameter sets that describe the experimental data for each model structure best, we performed numerical parameter estimation. The D2D framework is using a parallelized implementation of the CVODES ODE solver. The procedure of parameter estimation is based on multiple local optimizations for different initial guesses of the parameters. For the optimization, the LSQNONLIN algorithm (MATLAB, R2011a, Mathworks) was used. Most kinetic parameters were limited to values between 10^−6^ and 1. Exceptions include translocation parameters. Here, the upper boundary was raised to 10^2^. Parameter values close to upper or lower boundaries result from practical non-identifiability of the model structure. We assume that six orders of magnitude as a parameter range is sufficient to not hinder the parameter estimation process. For the random sampling of the multiple starting points, a Latin hypercube method was utilized. In addition to kinetic parameters, the observation function relating the ODE model to the experimentally accessible data contains scaling and noise parameters. These non-kinetic parameters were fitted in parallel to the kinetic parameters as described [53]. Using a previously established strategy [53], we ensured reliable convergence of our parameter estimation procedure for the two mathematical models (S2C Fig).

### Prediction profiles

To obtain confidence intervals of the model predictions for the additional internal feedback loops, we calculated predictions profiles for the respective species as described [54]. For our analysis, prediction profiles have been calculated along the complete time course of the core model with an additional intracellular feedback and species “internal_x_factor_mrna” (Fig 4B). Through the calculation of prediction profiles, a range for the specified trajectories of the species dynamic is given for each calculated time point, in which the likelihood value of the model stays within a 95% confidence level.

### Rankings (AIC/LRT)

Performances of different model structures are determined by the likelihood *L*. For comparison of the model structures, two different criteria are used (S2D,E Fig). First, we introduced a variation of the likelihood-ratio test:

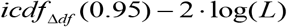

where *Δdf* denotes the difference in degrees of freedom between the two selected models and *icdf* denotes the inverse cumulative density function of the chi-squared distribution. The results of the likelihood ratio tests with the full model are then used to obtain the ranking of the corresponding model structures.

For the second criterion, all models are compared utilizing the Akaike Information Criterion (AIC), defined as:

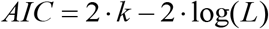

where *k* denotes the degrees of freedom in the respective model. While the AIC provides a ranking where each model is treated equally, the LRT provides information in terms of significance for a pairwise comparison of two selected models. In practice, the AIC slightly favors larger models due to the linear penalization of the degrees of freedom of a model.

## Data availability

Mircoarray data is available at the Geo database (https://www.ncbi.nlm.nih.gov/geo/query/acc.cgi?acc=GSE100928). All other relevant data are within the manuscript and its Supporting Information files.

## Acknowledgments

We thank Nao Iwamoto and Artyom Vlasov (German Cancer Research Center) for fruitful discussions and excellent technical assistance.

## Author contributions

KR generated the time–resolved quantitative gene expression data and performed all the experiments. AS generated the time–resolved quantitative signaling proteins data and the time–resolved quantitative gene expression data for the microarray. TM, JT and MS developed the mathematical model. CK analyzed the microarray data. JW, RB and MB provided the HCV system and contributed to the corresponding analyses. CE, JH, PAL and JGB performed the mouse experiments. XH performed single cell experiments. DS and GD provided the primary human hepatocytes. UK, JT, KR, AS, SC, FS, MT and MS conceived the project and wrote the manuscript. All authors contributed to and approved the paper.

## Supporting information captions

**S1 Fig. Core mathematical model of IFNα-induced JAK/STAT signaling and gene expression.** (**A**) Schematic representation of the core model according to Systems Biology Graphical Notation. TFBS: transcription factor-binding site. (**B**,**C**) Trajectories of the core model are shown together with the dynamic behavior of the core components of the JAK/STAT signaling pathway measured by quantitative immunoblotting (**B**) and to the expression of IFNα-induced genes examined by qRT-PCR (**C**) after stimulation of Huh7.5 cells with 500 U/ml IFNα. Filled circles: experimental data; line: model trajectories, shades: estimated error; a.u. arbitrary units.

**S2 Fig. The core model with an additional intracellular feedback is superior and IRF-downregulation enhances gene expression**. (**A**) Model rankings according to likelihood ratio test presented by the negative logarithmic likelihood penalized by parameter difference. Lower values indicate (**B**) Model rankings according to Akaike information criteria (AIC). The preferred model is the one with the smaller AIC value. (**C**) Assessment of the optimization performance by a waterfall plot. The best parameters were reproducibly found, which validates the applied model calibration approach. (**D**) Huh7.5 cells were growth factor depleted and pre-incubated for 24 hours with siRNA directed against *IRF2*, *IRF4* or *IRF8* or a combination thereof followed by 500 U/ml IFNa treatment. At indicated time points RNA was extracted and analyzed using qRT-PCR. Error bars represent SD (n=3). (**E**) Huh7.5 cells were growth factor depleted and pre-incubated for 24 hours with siRNA directed against IRF2 followed by 500 U/ml IFNa treatment. At indicated time points RNA was extracted and analyzed using qRT-PCR. Error bars represent SD (n=3).

**S3 Fig. IFNα-induced gene expression after co-stimulation with IL8 and the activation of STAT3 by IL1β is blocked by Stattic.** (**A**) Co-stimulation with IFNα and IL8. Huh7.5 cells were growth factor depleted followed by single treatment with 500 U/ml IFNα alone or in combination with 10 ng/ml IL8. At indicated time points RNA was extracted and analyzed using qRT-PCR. Error bars represent SD of biological triplicates. (**B**) Huh7.5 cells were single or co-stimulated with 500 U/ml IFNα and 10 ng/ml IL1β. Cells were lysed at indicated time points and analyzed using quantitative immunoblotting. Error bars represent SEM of three biological replicates. (**C**) mKate2-Stat3 primary mouse hepatocytes were stimulated with 5 ng/ml IL-6 or 10 ng/ml IL-1β. Line plots represent the dynamics of nuclear STAT3 for the indicated conditions. Two biological replicates and two technical replicates were included. Data represent mean ± SEM. (**D**) Huh7.5 cells were treated with 10 μM Stattic for up to 24 h or left untreated and cell viability was measured. (**E**) Huh7.5 cells were pre-treated with 10 μM Stattic followed by 10 ng/ml IL1β and 500 U/ml IFNα treatment. Cells were lysed at indicated time points and analyzed using quantitative immunoblotting. Error bars represent SD of biological triplicates. a.u.: arbitrary units.

**S4 Fig. Enhanced viral clearance in a HCV replicon cell line upon co-stimulation with IFNα and IL1β**. (**A**) Luciferase activity measurement in single and co-stimulated cells. Time-resolved measurements of luciferase activity in cells treated with 500 U/ml IFNα alone or in combination with 10 ng/ml IL1β compared to the unstimulated control. Error bars represent SEM of three biological replicates. (**B**) Luciferase activity measurement in single and co-stimulated cells. Time-resolved measurements of luciferase activity in cells treated with 50 U/ml IFNα alone or pre-treatment with 50 U/ml IFNα followed by 1 ng/ml IL1β treatment compared to the unstimulated control. Error bars represent SEM of four biological replicates.

**S5 Fig. Reduction of anti-viral T cell immunity following LCMV infection in IL1R1^−/−^ mice.** (**A**) Wild-type (WT) or IL1R1 knock-out (IL1R1^−/−^) CL57BL/6 mice were infected with 2×10^6^ pfu of LCMV WE. Four and eight days post infection, single cell suspensions from spleen and liver tissue as well as peripheral blood lymphocytes were stained using gp33 or np396 MHC class I tetramers or gp61 MHC II tetramer followed by staining with anti-CD8. Differences between WT and IL1R1^−/−^ cells were tested by two-way ANOVA. ***, p<0.001; **, p<0.01; *, p<0.05; n.s., not significant, n=6. (**B**) Four and eight days post infection, suspended liver cells or splenocytes were stained with the LCMV-specific peptides gp33, np396, or gp61. Additionally, surface staining with anti-CD8 or anti-CD4 antibodies and intracellular staining with anti-IFNγ antibodies was performed. Differences between WT and IL1R1^−/−^ cells were tested by two-way ANOVA. *, p<0.05; n.s., not significant, n=6. (**C**) Four and eight days post infection, lymphocytes were stained with antibodies against surface molecules.

**S1 Table. Initial concentrations of model species.** Measured concentrations (JAK1, TYK2, STAT1, STAT2, IRF9) were transformed from molecules per cell to nM by using STAT1 concentration as reference. Concentrations for receptors were assumed to be non-limiting and therefore set to a high amount [4].

**S2 Table. qRT-PCR primers and corresponding UPL probes**.

